# Mind the gap: quantifying individual–population gap in depressive symptom dynamics through energy landscapes

**DOI:** 10.64898/2026.05.20.726729

**Authors:** Masato Tsutsumi, Taisei Kubo, Takahiro A. Kato, Honda Naoki

## Abstract

People do not always feel as they appear. Someone who seems stable may struggle internally, whereas someone who appears distressed may experience it differently. This gap matters in psychiatry, where assessment relies on symptom scales and external evaluation. Here we developed mindGAP (Measuring INDividual-population GAPs in psychiatric energy landscapes), a hierarchical variational Bayesian framework that uses longitudinal questionnaire data to estimate population-level and individual-level pairwise maximum entropy models (pMEMs). We applied mindGAP to time-series PHQ-9 data from 248 participants. The population landscape contained three major states, whereas individual-level landscapes often diverged from this shared structure. We quantified this gap as individual-population landscape divergence, which was associated not only with depressive severity but also with modern-type depression-related traits (TACS-22) and interpersonal sensitivity-self traits (IPS-22). Thus, mindGAP opens a route to quantifying a previously unquantified gap between population-level and individual-level symptom organization.

## Introduction

Human mental states can be described from two distinct perspectives. At the population level, they are interpreted through externally observable patterns and population-derived assessment frameworks.

At the individual level, they are experienced subjectively. These two perspectives do not always coincide. An individual perceived as “cheerful” or “stable” may internally suffer from substantial distress, whereas someone regarded as “depressed” or “unwell” may subjectively feel relatively stable. Although such mismatches are intuitively familiar, they have rarely been formalized as a quantitative problem in a way that allows direct comparison between population-level evaluation and individual-level mental-state organization, even though related discrepancies between self-report and external assessment have long been recognized in psychiatry [1, 2].

Quantifying this gap requires measurement frameworks that can capture both population-level and individual-level symptom organization. In psychological and psychiatric research, mental states are commonly quantified using questionnaire scores, clinical ratings, and behavioral measures. These tools play a central role in assessment and clinical decision-making. In most cases, multiple items are summed or averaged into a small number of scalar indices, on the basis of which individuals are evaluated and compared. Although such representations are effective for identifying symptom severity and clinical risk, they compress complex symptom configurations into a limited set of summary measures, thereby obscuring heterogeneity in how symptoms are organized within individuals [3, 4]. As a result, individuals with similar scores may be treated as comparable even when their underlying symptom organization differs substantially. This makes it difficult to detect mismatches between population-based assessment and the way symptoms are organized within individuals.

To quantify such discrepancies in symptom presentation, mental states must be represented not merely as a small number of scalar indices but as a dynamical system that captures transitions among states over time. In psychiatric terms, states can be conceptualized as attractors, that is, configurations toward which an individual’s symptoms tend to maintain or return to following minor fluctuations. To characterize the stability and transitions among such attractors, a data-driven method called energy landscape analysis was originally developed to characterize attractors of brain activities [5–8]. This approach has recently been applied to the analysis of clinical and longitudinal mental health data [9, 10]. However, most existing studies estimate a single landscape from pooled data across individuals. As a result, it remains unclear whether population-level structures faithfully reflect the dynamics of individual patients, or whether substantial differences exist between collective and individual organizations.

In this study, we developed mindGAP (Measuring INDividual–population GAPs in psychiatric energy landscapes), a hierarchical variational Bayesian framework that jointly estimates population-level and individual-level symptom landscapes from longitudinal questionnaire data. Using longitudinal PHQ-9 data, we quantified individual–population landscape divergence, compared how the same observed symptom trajectories were mapped onto the two landscapes, and showed that individual-level features inferred solely from PHQ-9 responses were associated with and predicted scores on other psychological scales, supporting the validity of the inferred individual landscapes. Together, these results establish mindGAP as a quantitative framework for determining when population-level structure captures, or mischaracterizes, individual mental-state organization.

## Results

### Hierarchical landscapes link population-level and self-reported depressive states

We sought a unified framework capable of jointly representing individual structure of mental-state variability and its population-level structure (Fig. 1). We analyzed longitudinal self-reported responses to a standardized questionnaire assessing depressive tendencies (Fig. 1A). The data consist of high-dimensional categorical response patterns, which are binarized in preprocessing (see Methods).

**Fig. 1.**
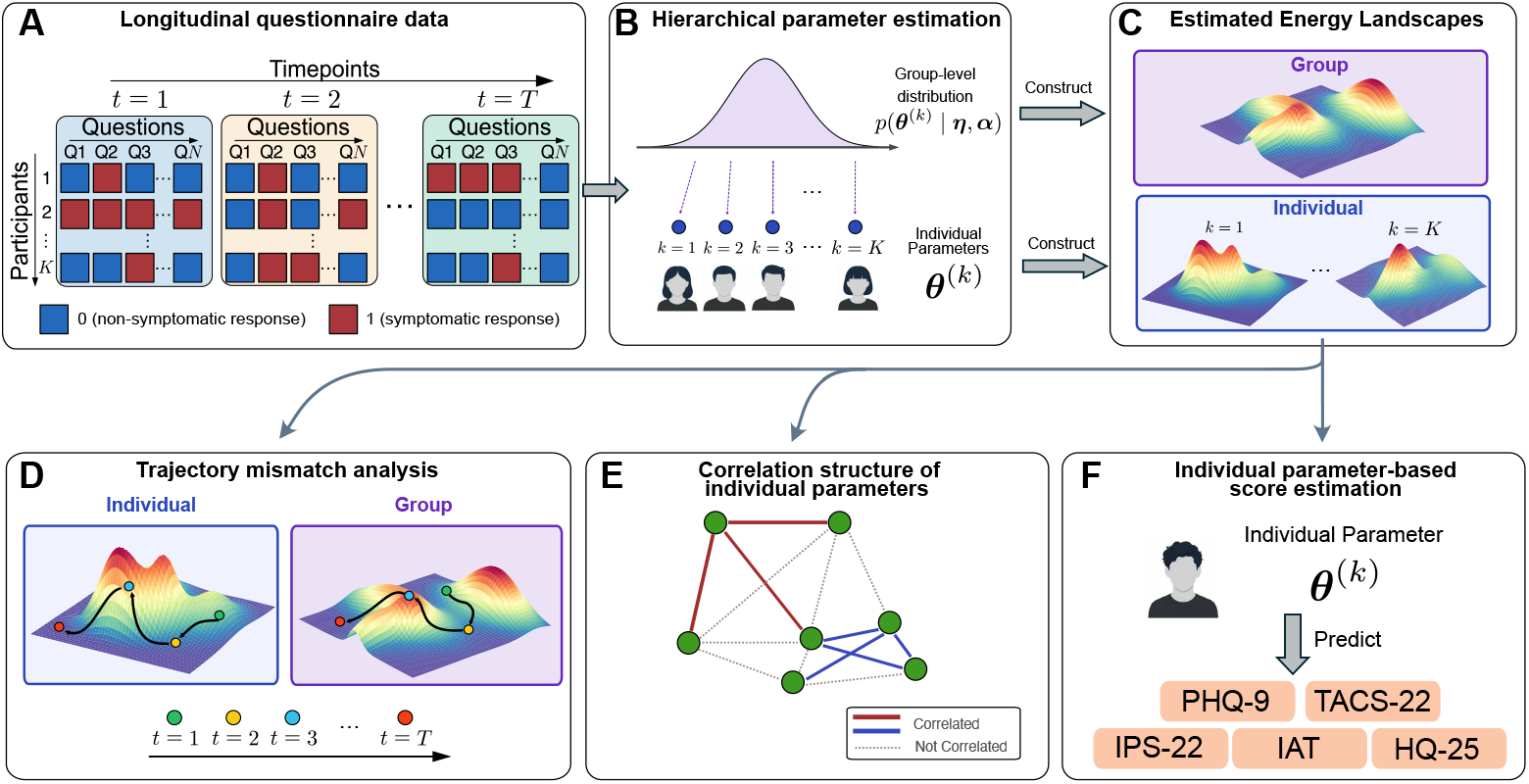
**A**, Longitudinal questionnaire data structure. *N* = 9 item responses of PHQ-9 were collected from *K* = 248 participants across *T* = 5 time points and binarized into symptomatic and non-symptomatic states, yielding a time series of multivariate symptom-response patterns. **B**, Each participant’s parameters are assumed to be drawn from a shared population distribution; the individual parameters are estimated jointly. **C**, Using the jointly inferred parameters, mindGAP reconstructs population-level and individual-level landscapes. **D**, Mapping of observed symptom trajectories onto individual-level and population-level landscapes to quantify mismatch in state organization. **E**, Cross-participant correlation structure of estimated interaction parameters, highlighting interactions that covary across individuals and those that vary more independently. **F**, Prediction of psychological assessment scale scores from individual parameters.

We represented binary PHQ-9 response patterns using a pairwise maximum entropy model (pMEM), equivalently an Ising-type Boltzmann distribution:

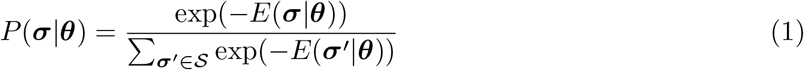

where ***σ*** represents a nine-bit vector consisting of the binarized questionnaire responses. The energy function *E*(***σ***) is defined as

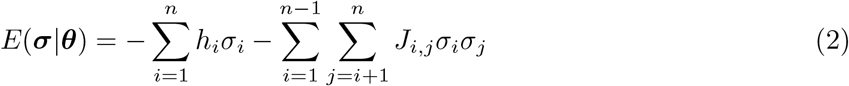

where *σ*_*i*_ denotes the *i*-th component of *bmσ*; *h*_*i*_ represents the baseline tendency of item *i* to take the value 1, and *J*_*i,j*_ represents the association between items *i* and *j*. In this context, “energy” serves as a quantitative measure of how likely a particular configuration of responses is: frequently observed response patterns are assigned lower energy values, whereas rarely observed configurations are assigned higher energy values.

mindGAP extends this pMEM representation by placing the individual parameter vectors 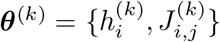 in a hierarchical model. For each bias and interaction parameter, values are allowed to differ across participants but are assumed to be drawn from a common population-level distribution with mean ***η*** and precision ***α*** (Fig. 1B). The mean vector ***η*** defines the population landscape, whereas each individual posterior mean ***µ***^(*k*)^ defines the individual-level landscape for participant *k*. These parameters were estimated using a variational expectation-maximization algorithm (see Methods) [11].

The following analyses proceeded in four stages. First, we constructed population-level and individual-level landscapes from the parameters estimated by mindGAP and characterized their stable states, or attractors, and basin structures (Fig. 1C). Second, we quantified population–individual mismatch by comparing how the same observed symptom trajectories were mapped onto the two landscapes (Fig. 1D). Third, we examined cross-participant variation in individual-level parameters, particularly the pairwise symptom interactions *J*_*i,j*_, to identify interaction structures underlying individual differences in symptom organization (Fig. 1E). Fourth, we tested whether individual-level parameters and individual–population landscape divergence were associated with, and could predict, scores on other psychological assessment scales (Fig. 1F).

### Observed score distributions and trajectories

Depressive tendencies were assessed using the Patient Health Questionnaire-9 (PHQ-9) [12], a widely used self-report questionnaire consisting of nine items related to depressive experiences and behaviors (Fig. 2A). Data were drawn from a longitudinal internet-based survey conducted in Japan, with repeated measurements collected across five time points. Each PHQ-9 item is scored on a scale from 0 (“not at all”) to 3 (“nearly every day”). In standard practice, responses to the nine items are summed to produce a total score ranging from 0 to 27 (Fig. 2B). Response distributions varied across items (Fig. 2C), indicating that similar total scores could arise from different combinations of item-level responses. We also examined whether clearly separated clusters could be identified in a low-dimensional representation of the item-level responses, but principal component analysis did not reveal such clusters (Fig. 2D). Across the five time points, most participants showed relatively stable total-score trajectories, whereas a subset exhibited marked fluctuations or persistently elevated values (Fig. 2E). Together, these observations motivated us to develop a method that can extract discrete response patterns from multivariate item-level data and characterize how those patterns change over time.

**Fig. 2.**
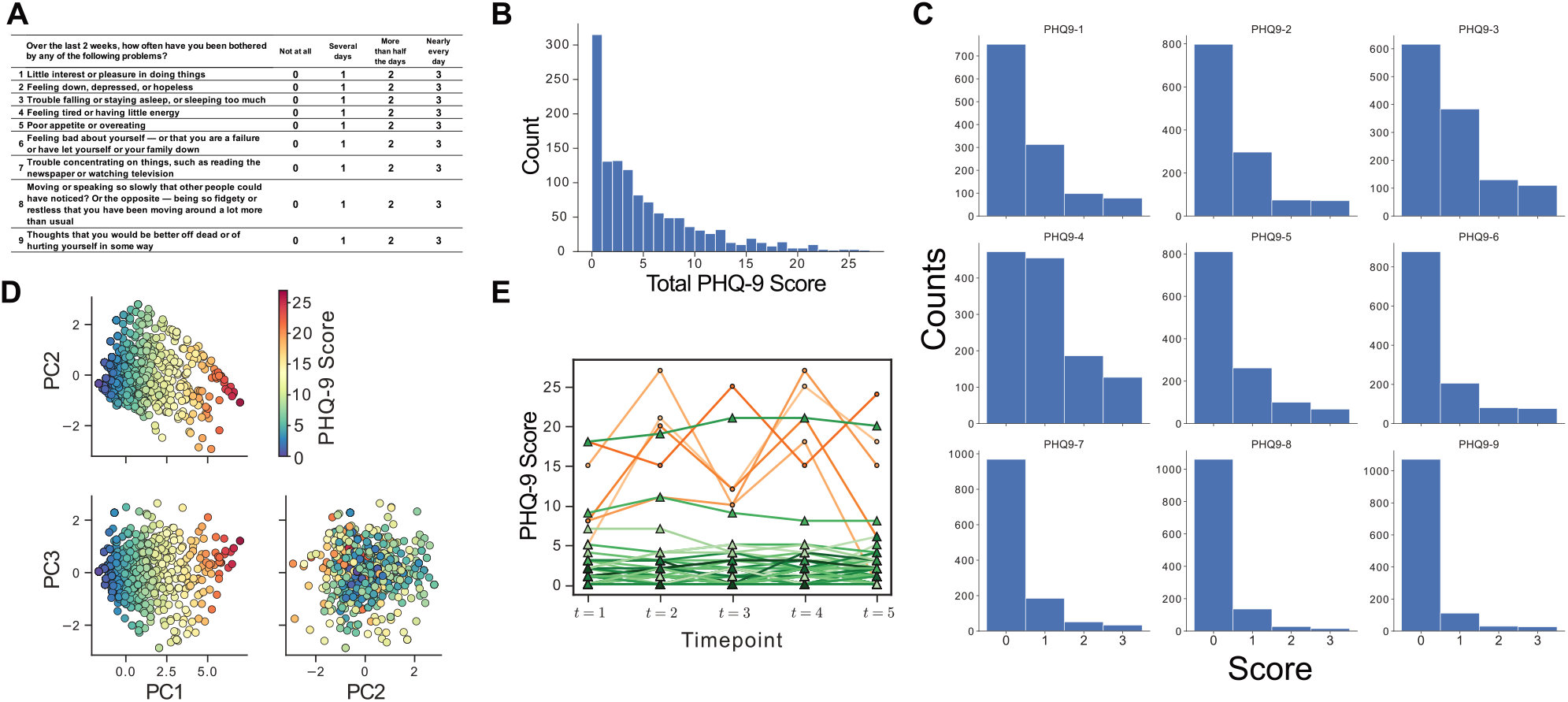
**A**, PHQ-9 questionnaire used in this study. The scale consists of nine items, each rated from 0 to 3 according to symptom frequency over the past two weeks. **B**, Distribution of total PHQ-9 scores across all participants and time points. **C**, Score distributions for each PHQ-9 item. **D**, Principal component analysis of item-level PHQ-9 responses. Each point represents one observation, colored by total PHQ-9 score. **E**, Individual PHQ-9 total-score trajectories across the five assessment waves. While the trend was generally stable at low or high levels (green), some cases showed a pattern of significant fluctuation (orange).

### Population-level energy landscape

Using mindGAP, we estimated the full set of parameters from the longitudinal PHQ-9 item-response data described above. We first focused on the population-level landscape determined by the population parameter vector ***η*** (Fig. 1 B,C). We identified local energy minima as attractors and defined each basin of attraction as the set of response patterns that converge to the same minimum through successive transitions to lower-energy neighboring states.

We identified three primary states (Fig. 3A): State 1, a resilient or asymptomatic state with minimal symptom endorsement across all items; State 2, an intermediate state, with symptoms present across PHQ-9 items 1–6 but absent across PHQ9 items 7-9; and State 3, a severe depressive state with persistent symptom endorsement across all items. State 1 was the most prevalent state, whereas State 2 and State 3 were much less common (0.71, 0.15, and 0.13, respectively; Fig. 3B). The transition matrix (Fig. 3C) showed high self-transition probabilities (0.97, 0.81, and 0.88) and stronger flow from State 2 to State 1 (0.11) than the reverse (0.025), consistent with the episodic nature of depression, in which many individuals improve whereas a smaller subgroup shows a more persistent or recurrent course [13].

**Fig. 3.**
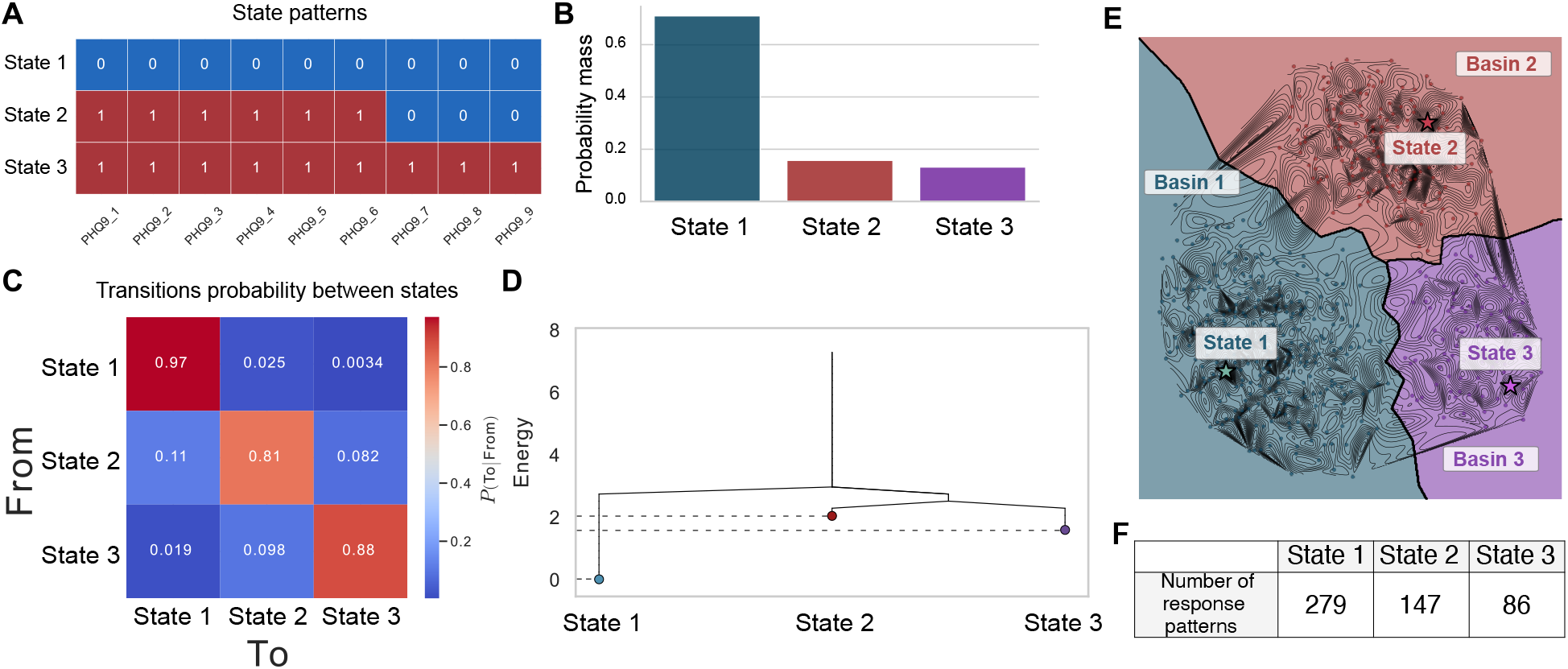
**A**, States identified in the population-level landscape, shown as binary response patterns across the nine PHQ-9 items. State 1 corresponds to minimal symptom endorsement, State 2 to an intermediate pattern, and State 3 to persistent symptom endorsement across items. **B**, Probability mass of each state, showing that State 1 occupies the largest proportion of the landscape. **C**, Transition probability matrix between states. **D**, Disconnectivity graph summarizing the energy barrier structure among the three states. The vertical axis indicates energy, and branch heights reflect the barriers separating attractors. **E**, Two-dimensional visualization of the population-level landscape. Colored regions indicate basins of attraction, black contour lines indicate interpolated energy levels, thick black lines indicate basin boundaries, and stars mark local minima corresponding to State 1–State 3. **F**, Number of binary response patterns assigned to each state.

To further assess the stability of each state and the accessibility of transitions between them, we quantified both state energies and the energy barriers separating states, and summarized the resulting topology as a disconnectivity graph (Fig. 3D). This analysis placed State 1 at the lowest energy and State 2 at the highest, with State 2 functioning as a metastable intermediate between the resilient and severe depressive states. Thus, State 1 appears to represent the most stable configuration, State 2 an unstable intermediate, and State 3 a relatively stable depressive configuration. To further illustrate the geometry of this landscape, we projected it into two dimensions for visualization (see Methods). The resulting landscape (Fig. 3E) revealed three basin regions, with a particularly broad basin around State 1. Although State 1 and State 3 were adjacent in this projection, direct transitions between them were rare, consistent with the low transition rate between these states. Notably, State 1 encompassed a substantially larger number of response patterns (*n* = 279) than State 2 (*n* = 147) or State 3 (*n* = 86), indicating greater configurational diversity within the resilient state. By contrast, State 2 and State 3 occupied a narrower range of patterns, suggesting more stereotyped symptom configurations, in line with previous reports in major depressive disorder [14–16].

### Individual-level energy landscape

Having characterized the population-level landscape, we next turned to the central question of whether these collective dynamics reflect the transitions experienced by individual participants. To address this, we constructed individual-level landscapes from posterior mean parameters ***µ***^(*k*)^ and examined how attractor transitions corresponded between the individual and population landscapes. As representative examples, trajectories of four participants were projected onto individual-level landscapes (Figs. 4A–D) and the population landscape (Fig. 4E). We identified four distinct patterns of individual-population correspondence: (a) steady–steady, in which the participant remained in the same attractor in both the individual and population landscapes (Fig. 4A); (b) steady–unsteady, in which the participant remained in the same attractor in the individual landscape but transitioned between different attractors in the population landscape (Fig. 4B); (c) unsteady–steady, in which the participant transitioned between different attractors in the individual landscape while remaining in the same attractor in the population landscape (Fig. 4C); and (d) unsteady–unsteady, in which the participant transitioned between different attractors in both landscapes (Fig. 4D). These contrasting patterns demonstrate that mismatches between population-level and individual-level evaluations are common, and that mindGAP can explicitly quantify such divergences in attractor transitions.

**Fig. 4.**
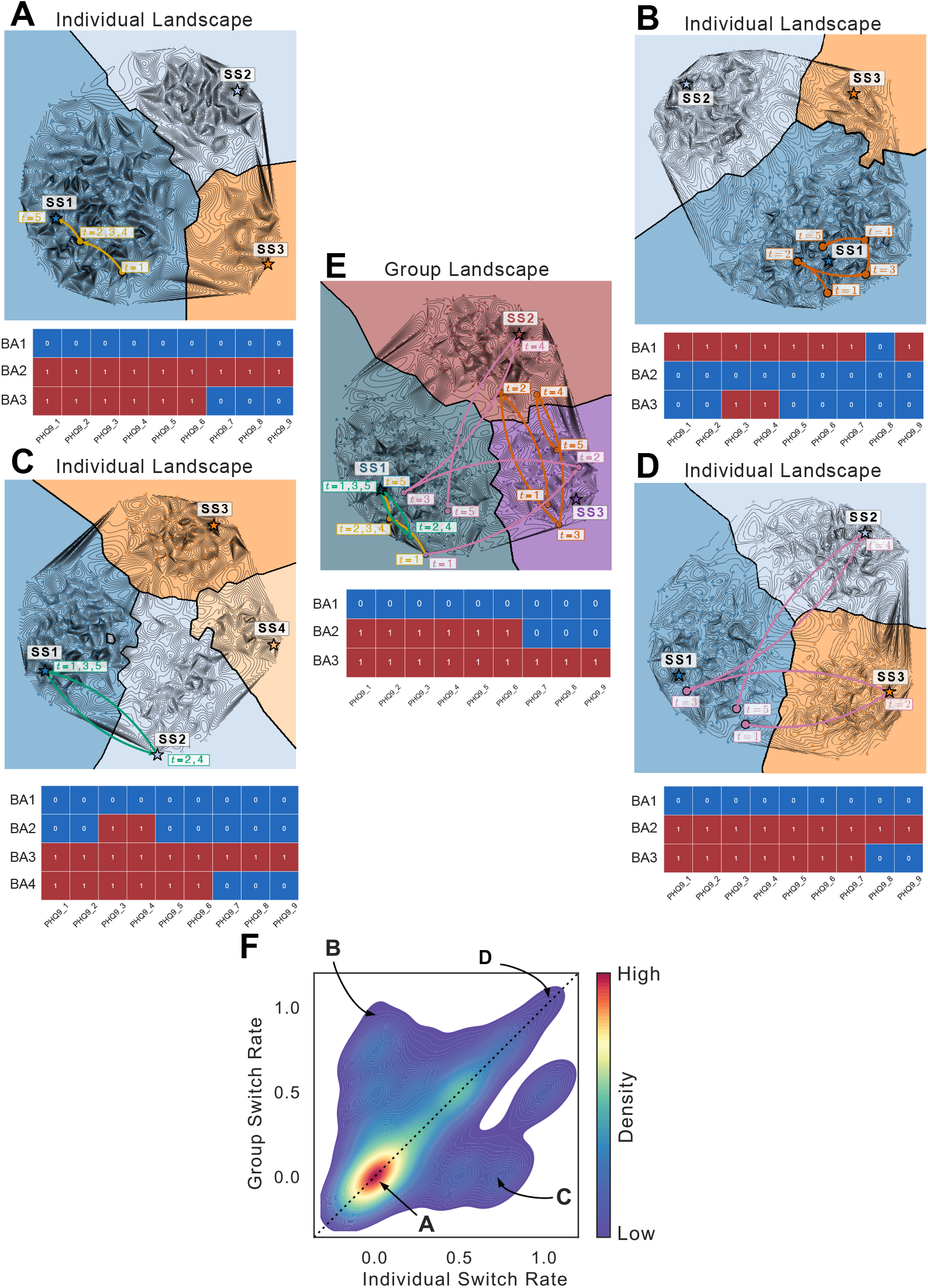
**A**–**D**, Four representative participants whose identical PHQ-9 response sequences are organized differently under individual-level and population-level landscapes. Coloured trajectories indicate observed transitions across five assessment waves; tables below show the binary PHQ-9 response patterns associated with each attractor state. **A** steady–unsteady: the trajectory remains within a single attractor in the individual landscape but switches attractors in the population landscape., **B**, unsteady–unsteady: the trajectory switches attractors in both the individual and population landscapes. **C**,steady–steady: the trajectory remains within the same attractor in both the individual and population landscapes. **D**, unsteady–steady: the trajectory switches attractors in the individual landscape but remains within a single attractor in the population landscape. **E**, The same four trajectories projected onto the population-level landscape, allowing direct comparison between individual-level and population-level interpretations of the same observed symptom sequences. **F**, Joint density plot of individual and population switching rates across all participants. Arrows **A**–**D** indicate regions corresponding to the four representative patterns shown in **A**–**D**.

To systematically compare these mismatches, we counted the number of basin transitions under both the individual-level and population-level landscapes and visualized the results as a density plot (Fig. 4F). This analysis showed that most participants remained in the same basin or changed basins only rarely in both landscapes (arrow A in Fig. 4F), whereas others showed mismatched transition frequencies between the two landscapes, corresponding to the unstable–stable and stable–unstable patterns (arrows B and C in Fig. 4F).

### A distinct interaction among PHQ-9 items 7–9 governs individual symptom dynamics

Because mindGAP estimates a full set of individual-level parameters for each participant, we next asked how individual differences in these parameters were organized across the population. Each individual-level parameter vector consists of item-wise bias terms *h*_*i*_, which represent the baseline propensity of each symptom item to be endorsed, and pairwise interaction terms *J*_*i,j*_, which represent the coupling between pairs of symptoms. We therefore examined the correlation structure of the estimated *h*_*i*_ and *J*_*i,j*_ parameters across participants.

The *h*_*i*_ parameters showed no clear correlation structure across items (Fig. 5A), indicating that individual differences were not primarily organized as a shared tendency to endorse a specific subset of PHQ-9 items. In contrast, the *J*_*i,j*_ parameters exhibited a marked correlation structure (Fig. 5B). In particular, the *J*_*i,j*_ correlation matrix showed coherent blocks corresponding to interaction modules defined in Fig. 5C. Interactions between item 7 and items 1–6 were strongly correlated with one another within the S7 module, indicating that participants with strong coupling between item 7 and one background symptom also tended to show strong coupling between item 7 and other background symptoms. Similar patterns were observed for the S8 and S9 modules. By contrast, direct interactions among items 7–9 (T module) showed moderate internal coherence, whereas background interactions among items 1–6 (BG module) showed comparatively weak and less coherent structure. Quantification of within-module mean absolute edge correlation confirmed that the S7, S8, S9 and T modules were more coherently organized than the BG module (Fig. 5D). These results suggest that PHQ-9 items 7–9 act as dominant interaction axes around which the coupling of items 1–6 is organized across individuals (Fig. 5E).

**Fig. 5.**
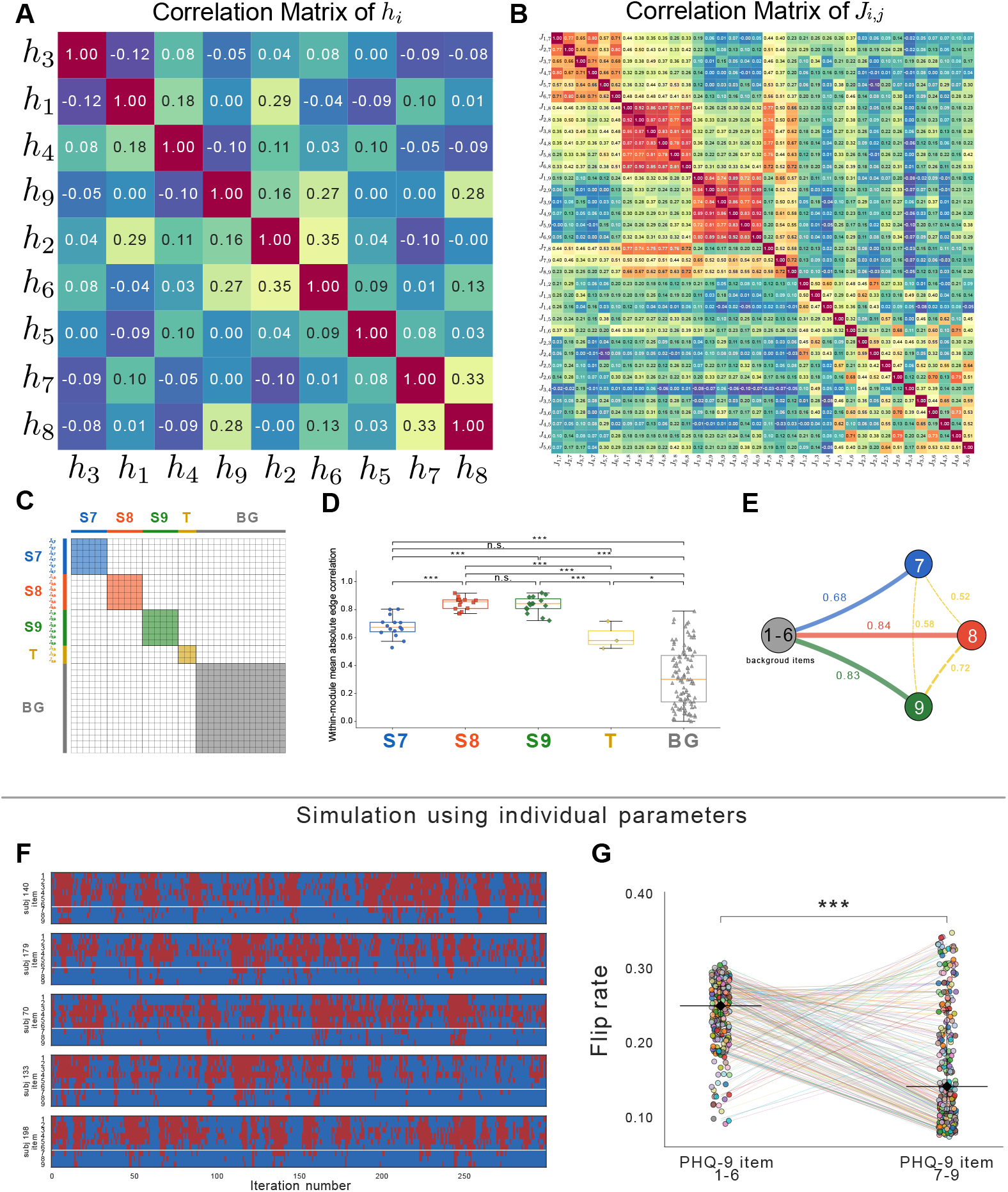
**A**, Across-participant correlation matrix of item-wise bias parameters *h*_*i*_. **B**, Across-participant correlation matrix of pairwise interaction parameters *J*_*i,j*_. **C**, Module schematic. Pairwise interactions were grouped into five modules: S7, S8, and S9, comprising interactions between item 7, 8, or 9 and background items 1–6; T, comprising direct interactions among items 7–9; and BG, comprising interactions among items 1–6. **D**, Within-module mean absolute edge correlation for each module across participants. **E**, Concept diagram summarizing mean absolute edge correlations among item groups 1–6, 7, 8 and 9. **F–G**, Simulation of questionnaire responses using each participant’s estimated parameters. **F**, Representative simulated sequences of binary questionnaire responses. **G**, Flip rates calculated from the simulated responses, compared between PHQ-9 items 1–6 and items 7–9. Lower values indicate less frequent response changes. P values are indicated as ***(*p <* 0.001), **(*p <* 0.01), *(*p <* 0.05), n.s., not significant.

To test whether this parameter-level organization was reflected in model-implied symptom dynamics, we next performed simulations using Markov chain Monte Carlo methods (MCMC). For each participant, we generated binary symptom-state trajectories by sampling from the Boltzmann distribution defined by that participant’s landscape. This allowed us to examine whether the inferred interaction structure produced a dynamical pattern consistent with the hypothesized hierarchical organization, in which items 7–9 behave as slowly switching variables and items 1–6 fluctuate in relation to them. Representative simulated trajectories showed that PHQ-9 items 7–9 tended to remain in the same binary state over longer stretches of sampling iterations, whereas items 1–6 switched more frequently (Fig. 5F). We quantified this tendency using the item-level flip rate, defined as the frequency of binary state switches between consecutive MCMC iterations. Compared with items 1– 6, PHQ-9 items 7–9 showed significantly lower flip rates (Fig. 5G), indicating that these items were more dynamically stable under individual-level landscapes.

### Individual-level landscape structure reflects broader psychological traits

To quantify how each individual landscape differed from the population landscape, we computed the Jensen–Shannon divergence (JSD) between their probability distributions. Larger values indicate greater divergence, which varied substantially across participants (Fig. 6A). We next examined whether this landscape distance was associated with scores on five psychological questionnaires available for all participants: PHQ-9 [12], IPS-22 [17], IAT [18], TACS-22 [19], and HQ-25 [20]. Greater landscape distance was positively associated with higher scores on all five scales (Fig. 6 B–G).

**Fig. 6.**
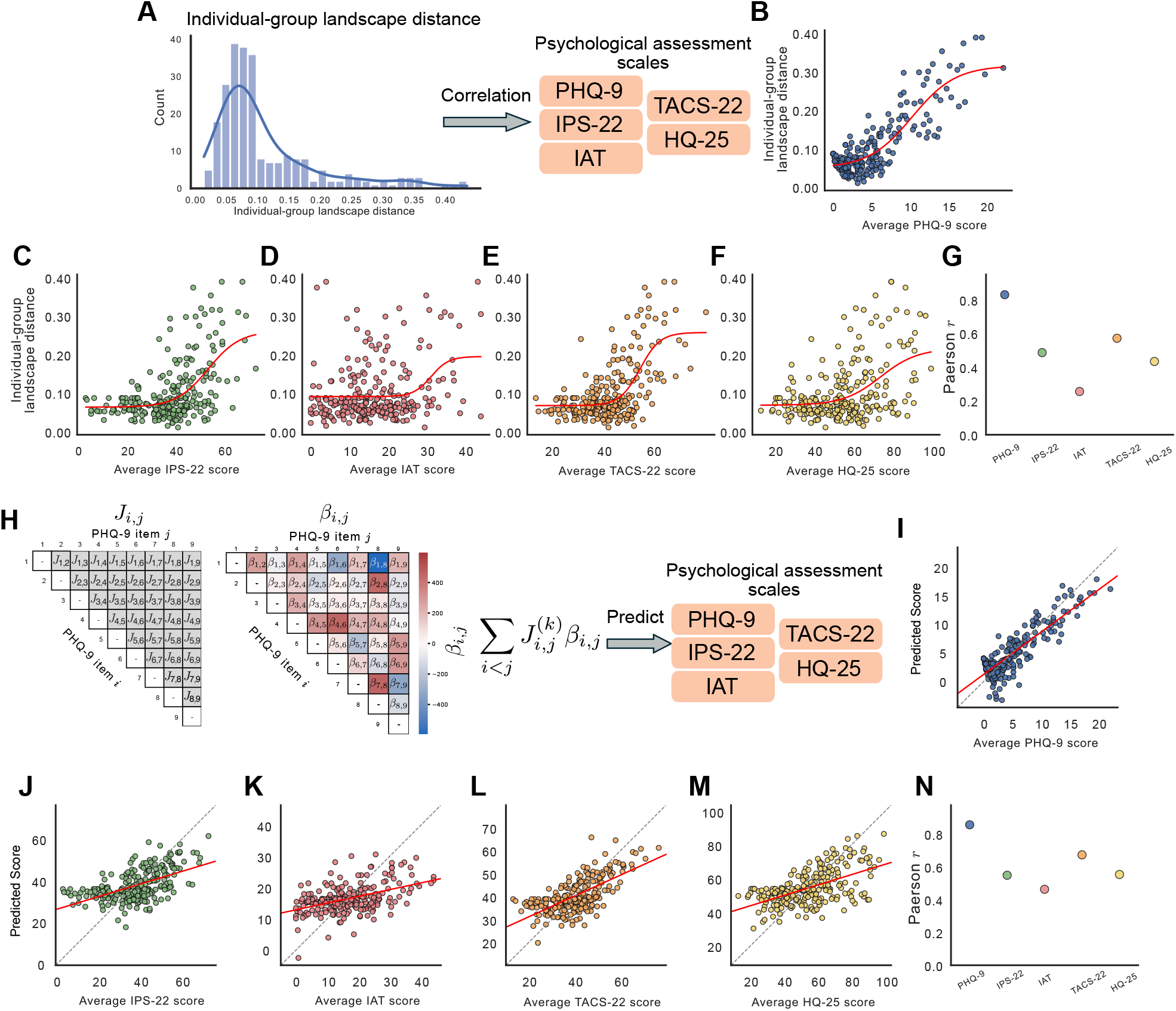
**A**, Distribution of individual–population landscape distance (JSD) across participants. **B**–**F**, Relationship between landscape distance and time-averaged scores on PHQ-9 (**B**), IPS-22 (**C**), IAT (**D**), TACS-22 (**E**) and HQ-25 (**F**). Each point represents one participant; red curves indicate fitted sigmoid functions. **G**, Pearson correlation between landscape distance and average questionnaire score for each scale. **H**, Schematic of score prediction from individual-level interaction parameters. Participant-specific 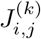 values were estimated from PHQ-9 data using mindGAP. For each questionnaire, scale-specific weights *β*_*ij*_ were fitted and evaluated by leave-one-out cross-validation. **I**–**M**, Leave-one-out predicted versus observed average scores for PHQ-9 (**I**), IPS-22 (**J**), IAT (**K**), TACS-22 (**L**) and HQ-25 (**M**). Red lines indicate linear fits. **N**, Pearson correlation between predicted and observed average scores for each scale.

We then asked whether the richer individual-level interaction structure could predict these questionnaire scores. Using the participant-specific pairwise interaction parameters 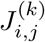 inferred from longitudinal PHQ-9 responses, we fitted scale-specific regression models and evaluated their predictive performance using leave-one-out cross-validation (Fig. 6H). Predicted scores were significantly correlated with observed average scores for all five questionnaires (Fig. 6I–N). For TACS-22, HQ-25, and IAT, prediction from the interaction parameters was comparable to or stronger than the corresponding association with landscape distance, indicating that individual interaction structure captures information not fully represented by the summary divergence measure. Together, these cross-questionnaire associations and predictions support the validity of the inferred individual-level landscapes despite the limited number of observations available for each participant.

## Discussion

In this study, we established mindGAP (Measuring INDividual–population GAPs in psychiatric energy landscapes), a hierarchical framework for estimating population-level and individual-level landscapes from longitudinal questionnaire data. This allowed us to directly compare population-level and individual-level symptom structures and address whether population-level structure adequately reflects individual dynamics. Applying mindGAP to longitudinal PHQ-9 data, we found that although the population-level landscape exhibited three major states, individual-level landscapes often differed substantially from this pattern. These discrepancies reveal a measurable gap between population-level and individual-level symptom organization—the central quantity captured by mindGAP.

The three population-level states were clinically interpretable: State 1 represented a low-symptom or asymptomatic state, State 2 an intermediate depressive state, and State 3 a globally symptomatic and severe depressive state. State 2 was characterized by endorsement of PHQ-9 items 1–6 but not items 7–9, indicating affective, somatic, and self-evaluative symptoms without concentration difficulty, psychomotor change, or suicidal ideation. In contrast, all nine PHQ-9 items were endorsed in State 3, including these cognitive, psychomotor, and suicidal symptoms. The separation between States 2 and 3 is consistent with the view that major depressive disorder is a heterogeneous syndrome rather than a unitary disease entity, and is reminiscent of long-standing clinical distinctions between melancholic or endogenous depression and broader non-melancholic depressive presentations [12, 14, 21–23]. However, these states should not be interpreted as formal diagnostic subtypes. Rather, they show that population-level landscapes estimated using mindGAP can capture clinically recognizable differences in depressive symptom organization from longitudinal item-level data.

Importantly, although inferred solely from longitudinal PHQ-9 responses, individual–population landscape distance was associated with scores on other psychological questionnaires, including TACS-22 and IPS-22 (Fig. 6 A–G). Moreover, participant-specific interaction parameters predicted scores on these questionnaires in leave-one-out cross-validation, with predictive associations comparable to or stronger than those obtained using landscape distance for several scales (Fig. 6H–N). These cross-questionnaire associations and predictions indicate that the inferred individual-level structure captures psychologically meaningful variation shared across assessment instruments. Because each participant’s parameters were estimated from only five time points, the precision of individual estimates may be limited. Nevertheless, their consistent relationships with other questionnaire measures provide convergent support for the validity of the inferred individual-level landscapes. Together, these findings highlight the utility of mindGAP for quantifying meaningful gaps between population-level and individual-level mental-state organization.

The present study can also be positioned in relation to recent mental-health applications of energy landscape analysis. For example, Tatematsu et al. applied this approach to longitudinal K6 data from Japanese high school students during the COVID-19 pandemic and demonstrated that psychological distress can be represented through temporal transitions among landscape states, including subgroup-specific structures [10]. Their study established the usefulness of energy landscapes for capturing heterogeneity beyond conventional summary scores. However, the analysis remained at the pooled or subgroup level. Here, mindGAP extends this framework to the individual level by estimating a distinct landscape for each participant and directly quantifying its divergence from the population landscape.

More broadly, the present study should also be viewed in the context of the wider energy-landscape literature, which originated in neuroscience. Watanabe and colleagues pioneered a data-driven framework for reconstructing energy landscapes from brain activity using pairwise maximum entropy models, first in resting-state fMRI and later in studies of perceptual dynamics and autism [5, 6, 24, 25]. Ezaki and colleagues subsequently systematized this methodological line of work and further expanded its application to EEG data and cognitive traits [7]. More recently, Watanabe and colleagues combined individual-level energy-landscape analysis with brain-state-driven TMS to test causal interventions on atypical brain dynamics and behavior [26]. However, even these person-specific studies did not aim to jointly estimate population- and individual-level landscapes within a unified framework and directly quantify their discrepancy. In this respect, the present study extends the landscape framework by treating divergence between individual and population landscapes as a primary inferential target.

Energy landscape analysis has also been directly applied to ecological data. Suzuki et al. showed that this framework can elucidate multistability in ecological communities across environmental gradients [27]. Relatedly, Toju et al. extended the broader landscape perspective in microbiome research by outlining stability-landscape-inspired assembly statistics for ecosystem-scale inference, forecasting, and management of abrupt community shifts [28]. These studies underscore the flexibility of landscape-based approaches across complex biological systems. However, such analyses often assume a fixed landscape, even though attractor structure may vary with environmental conditions or across regime shifts. mindGAP provides one possible extension of this line of work by enabling overall and condition-specific landscapes to be estimated separately but within a shared framework. This may be particularly useful for ecological data, where the landscape before and after a regime shift may not be identical, but may instead reorganize as the environment changes.

Overall, these findings highlight the promise of individual-level landscape estimation for precision psychiatry, particularly for individuals whose symptom organization may be poorly represented by population-average models. At the same time, the present study is limited by sparse longitudinal sampling, discrete state representations, and the use of a single cohort. Future studies incorporating denser longitudinal measurements, multimodal validation, and prospective prediction will be essential for determining whether individual-level landscape estimation can evolve from a descriptive framework into a practical tool for precision psychiatry.

## Methods

### Questionnaire data

This longitudinal observational study complied with the Declaration of Helsinki and was approved by the ethics committees of Kyushu University, Japan (approval no. 25-84). Data were collected via an online survey platform (Cross Marketing Inc., Tokyo, Japan). All participants provided informed consent electronically, participation was voluntary, and responses were anonymized before analysis. The broader survey project has also been used in previous publications.

Initial recruitment and quality-control procedures in the four-wave cohort were as follows: 2,182 individuals were invited to the screening survey, 1,509 responded, and eligibility criteria included age 30–59 years, Japanese language proficiency, and balanced sex/age strata (30s, 40s, 50s). Of these, 1,178 full-time employees (employed since at least June 2019) participated at baseline (June 2020). In the four-wave framework (June 2020, December 2020, August 2021, April 2022), 561 completed both baseline and wave 4; after excluding 107 invalid responders based on trap questions and 128 participants with pre-pandemic low outing frequency (≤3 days/week in December 2019), 326 remained. In the present study, we further extended follow-up to a fifth wave and analyzed participants with complete data across five time points: approximately every six months to one year from June 2020, immediately following the state of emergency declaration, through December 2020, August 2021, April 2022, and finally in August 2023. A total of *K* = 248 participants completed all five assessments and were included in the present analyses. The primary measure used in this study was the Patient Health Questionnaire-9 (PHQ-9) [12], a 9-item self-report questionnaire that assesses depressive symptoms over the past two weeks. Each item is rated on a 4-point scale (0 = not at all, 1 = several days, 2 = more than half the days, 3 = nearly every day).

For the energy landscape analysis, we binarized responses such that scores of 2 or 3 were coded as 1 (symptomatic) and scores of 0 or 1 were coded as 0 (non-symptomatic), following previous energy landscape studies of questionnaire data [10]. This binarization allows us to represent each response pattern as a binary state vector in the 2^9^ = 512 dimensional state space.

In addition to PHQ-9, participants also completed several other psychological assessment scales at each time point, including the Interpersonal Sensitivity Scale-22 (IPS-22) [17], the Internet Addiction Test (IAT) [18], the Test of Compulsive Internet Use Scale-22 (TACS-22) [19], and the Hikikomori Questionnaire-25 (HQ-25) [20]. These additional measures were used to evaluate associations between individual–population landscape distance and psychological constructs (Fig. 6A–G), and to test whether individual-level interaction parameters estimated from PHQ-9 data could predict external questionnaire scores using leave-one-out cross-validation (Fig. 6H–N), as described in the Results section.

### Boltzmann Representation of Questionnaire Response Patterns

We represented questionnaire response patterns using a pairwise maximum entropy model (pMEM), equivalently a Boltzmann distribution over all possible binary symptom states. Let

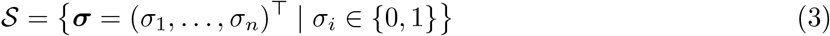

denote the state space of all possible response patterns for a questionnaire with *n* = 9 items. Here, *σ*_*i*_ = 1 indicates a symptomatic response for item *i* (i.e., scores of 2 or 3), whereas *σ*_*i*_ = 0 indicates a non-symptomatic response (i.e., scores of 0 or 1). At each time point *t*, the observed response pattern *σ*(*t*) is an element of S. Because *n* = 9, the state space contains 2^9^ = 512 possible response patterns.

We assumed that the probability of observing a response pattern ***σ*** follows a Boltzmann distribution,

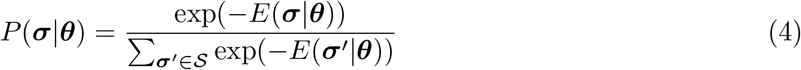

Under this formulation, response patterns with lower energy have higher probability. Accordingly, local minima in the energy landscape correspond to stable or recurrent symptom configurations. The energy function was parameterized as

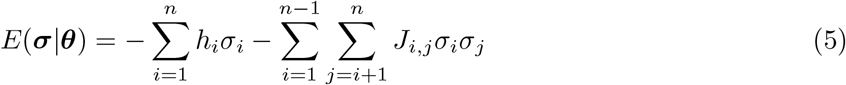

The parameter vector ***θ*** = (***h, J*** )^⊤^ characterizes the symptom response landscape. The parameter *h*_*i*_ represents the basal tendency toward a symptomatic response for item *i*, whereas *J*_*i,j*_ represents the pairwise interaction between items *i* and *j*. This Ising-type Boltzmann distribution is mathematically equivalent to a pairwise maximum entropy model (pMEM), namely, the maximum-entropy distribution constrained by the first- and second-order statistics of the binary questionnaire responses [5–11, 24–29].

### Likelihood of the Pairwise Maximum Entropy Model

To estimate the individual parameter vector, we next introduce the likelihood of the pairwise maximum entropy model for each participant. For participant *k*, let

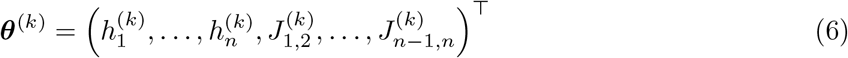

denote the individual parameter vector. Let

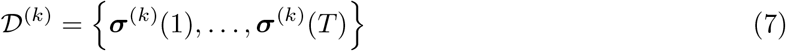

denote the observed response sequence for participant *k*, where ***σ***^(*k*)^(*t*) is the response pattern vector at time point *t*. The likelihood of the observed data for participant *k* is given by

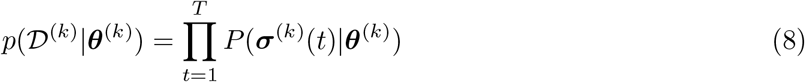

Here,

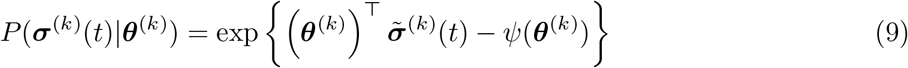

Where

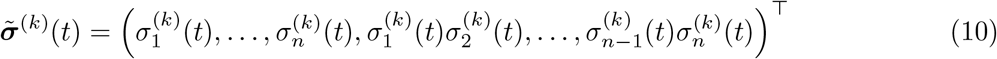

is the sufficient-statistic vector at time point *t*, and

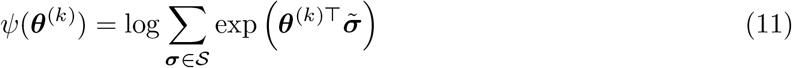

is the log-partition function. The corresponding log-likelihood is

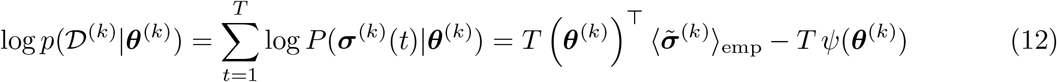

Where

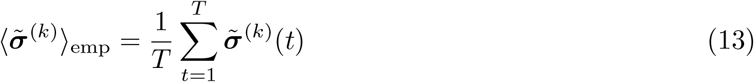

denotes the empirical mean of the sufficient-statistic vector for participant *k*.

In previous studies, parameters were typically estimated by maximizing this log-likelihood independently for each participant [5–7, 10, 11, 29]. In the present study, however, mindGAP incorporated this likelihood into a hierarchical variational Bayesian framework to jointly estimate individual-level and population-level parameters.

### mindGAP: Variational Bayesian Estimation of Individual and Population Parameters

mindGAP implements a hierarchical variational Bayesian model to estimate individual-level and population-level parameters jointly. To estimate a full pairwise maximum entropy model for each participant while sharing information across participants, we introduced a hierarchical Bayesian model. Specifically, the individual parameter vector ***θ***^(*k*)^ was assumed to arise from a population-level Gaussian distribution,

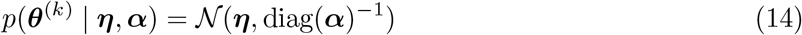

where ***η*** is the population-level mean parameter vector and ***α*** is the precision vector. Under this hierarchical model, ***η*** represents the average symptom landscape across participants, whereas ***θ***^(*k*)^ represents participant *k*’s deviation from that average.

Combining the likelihood defined in the previous section with this prior, the posterior distribution for participant *k* is given by

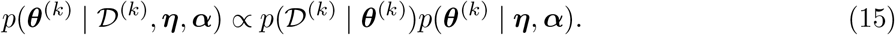

Because this posterior distribution is not available in closed form, we approximated it using a diagonal Gaussian variational distribution,

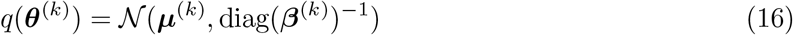

where ***µ***^(*k*)^ and ***β***^(*k*)^ denote the variational posterior mean and precision, respectively.

The variational distribution was estimated by maximizing the evidence lower bound (ELBO),

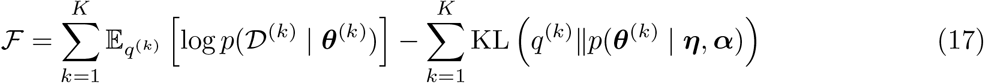

where KL(·∥·) denotes the Kullback–Leibler divergence. The first term measures how well the individual models explain the observed response patterns, whereas the second term regularizes the individual parameters toward the population-level distribution while accounting for uncertainty in the variational posterior.

The main computational difficulty in maximizing the ELBO lies in the log-partition function *ψ*(***θ***^(*k*)^) contained in the likelihood term. Following previous variational approaches for pairwise maximum entropy models [11, 29], we approximated *ψ*(***θ***^(*k*)^) by a second-order Taylor expansion around the current population mean ***η***,

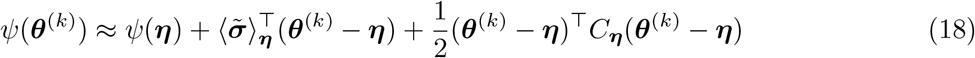

where 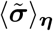 is the model-expected sufficient-statistic vector under ***η***, and *C*_***η***_ is its correlation matrix. Because the PHQ-9 response space contains only 2 = 512 possible binary states, both quantities were computed exactly by enumerating all states rather than by sampling.

### Variational Expectation-Maximization (VEM) Algorithm

We estimated the individual variational parameters and the population-level hyperparameters using a variational expectation-maximization algorithm. A step-by-step derivation of the resulting update equations is provided in the Supplementary Information. The algorithm alternates between two steps. In the E-step, the population-level parameters ***η*** and ***α*** are held fixed, and the variational posterior for each participant is updated according to the following closed-form expressions:

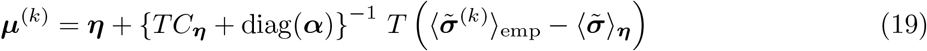

And

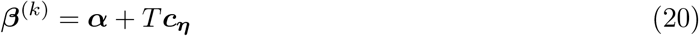

Where

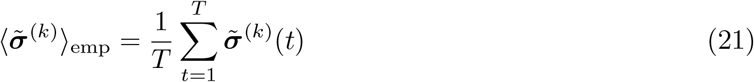

and ***c***_***η***_ denotes the vector of diagonal elements of *C*_***η***_

In the M-step, the individual variational posterior distributions are held fixed, and the population-level parameters are updated as

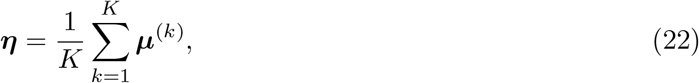

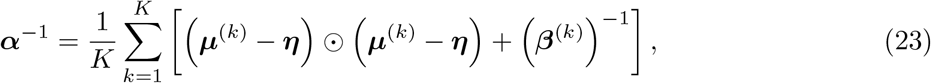

where ⊙ denotes element-wise multiplication.

The E-step and M-step were alternated until convergence. After convergence, the population-level landscape was constructed from ***η***, and each individual-level landscape was constructed from the corresponding posterior mean ***µ***^(*k*)^. In the main analysis, the five observations from each participant were used to estimate one individual-level parameter vector.

### Characterization of Energy Landscapes

To characterize the landscapes, we calculated the energies of all 2^*n*^ possible binary response patterns using Eq. (5). For the population-level and individual-level analyses, these energies were computed using ***η*** and ***µ***^(*k*)^, respectively.

#### Identification of attractors and basins

We identified attractor states using the following deterministic procedure. Starting from an arbitrary initial state, we examined all neighboring states reachable by a single-bit flip and moved to the one with the lowest energy if its energy was lower than that of the current state. This update was repeated until no neighboring state had lower energy. The final state reached by this procedure was defined as an attractor state. Repeating the same procedure for all possible initial states partitioned the state space into basins, where each basin consisted of the set of states that converged to the same attractor. The probability mass of each basin was then computed as the sum of the Boltzmann probabilities of all states belonging to that basin.

#### Basin-to-basin transitions and disconnectivity graph

We defined basin-to-basin barriers as the minimum, over paths on the Hamming graph, of the maximum energy along the path, measured from the starting attractor’s energy; the search used the algorithm of [8, 30]. The pairwise barriers were assembled into a disconnectivity graph.

#### Two-dimensional visualization of the energy landscape

To visualize the full energy landscape, we represented the state space as a weighted Hamming graph and embedded it in two dimensions using the Fruchterman-Reingold algorithm [31]. Each of the 2^*n*^ states was treated as a node, and edges were placed between neighboring states differing by a single-bit flip. Edge weights were set to 10 for state pairs within the same basin and to 0.5 for state pairs belonging to different basins, so that states within the same basin were placed closer together in the two-dimensional layout.

#### Interpolation of energy contours

Based on the two-dimensional node arrangement obtained by the Fruchterman-Reingold algorithm [31], we interpolated the energy values on a regular grid from the discrete node energies using a cubic interpolation algorithm [32–34]. Contour lines were then drawn from the resulting interpolated energy field.

### Quantification of Individual–Population Landscape Divergence

To quantify the divergence between individual-level and population-level landscapes, we computed the Jensen-Shannon Divergence (JSD):

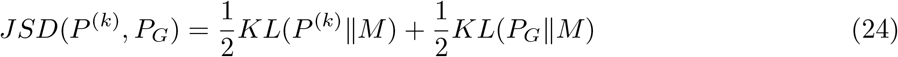

where *P* ^(*k*)^ is the probability distribution over all 2^*n*^ states for participant *k*’s, *P*_*G*_ is the population-level distribution, 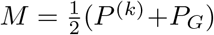 is the mixture distribution, and *KL* denotes the Kullback-Leibler divergence. Because JSD is symmetric (i.e., *JSD*(*P* ^(*k*)^, *P*_*G*_) = *JSD*(*P*_*G*_, *P* ^(*k*)^)), this metric quantifies how much an individual’s landscape deviates from the population-level landscape.

### Association between landscape distance and psychological assessment scores

Individual–population landscape distance was defined as the Jensen–Shannon divergence between the Boltzmann distributions of the individual-level landscape and the population-level landscape, as described above. For each participant and questionnaire, we computed the time-averaged total score across the five assessment waves. We examined the relationship between landscape distance and each average questionnaire score across participants (Fig. 6A–F). Nonlinear associations were summarized using sigmoid fits, and association strength was quantified using the Pearson correlation between landscape distance and average score for each scale (Fig. 6G).

### Prediction of psychological assessment scores from *J*_*i,j*_

To evaluate whether individual-level interaction structure predicted external questionnaire scores, we used individual pairwise interaction parameters 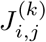 estimated from longitudinal PHQ-9 data using mindGAP as predictor variables. For each questionnaire, we fitted scale-specific regression weights *β*_*ij*_ that mapped these 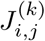 values to the participant’s time-averaged score:

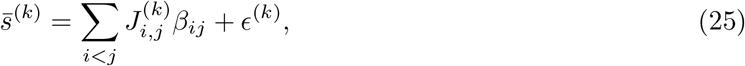

where 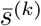 denotes the time-averaged score for participant *k* on a given questionnaire and *β*_*ij*_ denotes the scale-specific regression weights (Fig. 6H). Predictive performance was evaluated using leave-one-out cross-validation. For each held-out participant *k, β*_*ij*_ was estimated from the remaining *K* − 1 participants and used to compute the predicted score 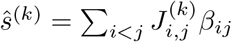. Performance was quantified as the Pearson correlation between leave-one-out predicted and observed average scores across all participants for each scale (Fig. 6I–N).

### Quantification of interaction-module organization

To examine how estimated *J*_*i,j*_ parameters were organized across participants, we grouped the 36 pairwise interactions into five modules (Fig. 5C): S7, S8 and S9 (interactions between item 7, 8 or 9 and background items 1–6), T (direct interactions among items 7–9, namely *J*_7,8_, *J*_7,9_ and *J*_8,9_) and BG (background interactions among items 1–6). For each module, we computed the mean absolute Pearson correlation among all within-module *J*_*i,j*_ pairs across participants. Module-level comparisons were displayed as box plots (Fig. 5D). For visualization of the broader organization among item groups, we also summarized mean absolute edge correlations among groups 1–6, 7, 8 and 9 in a concept diagram (Fig. 5E).

### MCMC-based assessment of item-level stability

To examine whether the target items showed distinct dynamical stability, we defined the background item set as ℐ _bg_ = 1, … , 6 and the target item set as ℐ _tar_ = 7, 8, 9.

For each participant, we generated binary response trajectories from the individual-level pMEM using Gibbs sampling. During each Gibbs sweep, the nine items were updated in random order. Conditional on the current states of the other items, item *i* was sampled according to

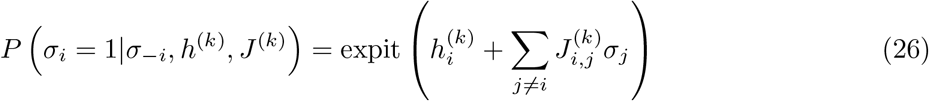

where *σ*_*i*_ ∈ 0, 1 denotes the current binary state of item *i, σ*_−*i*_ denotes the states of all items except item *i*, and expit(*x*) = 1*/*(1+exp(−*x*)). Thus, the probability that item *i* was sampled as symptomatic depended on its individual bias and on the summed influence of the other currently active items.

For each participant, the first *B* = 5,000 sweeps were discarded as burn-in, and the subsequent *M* = 10,000 sweeps were retained for analysis. Let 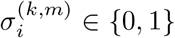 denote the retained MCMC state of item *i* for participant *k* at saved sweep *m* = 1, … , *M* .

For visualization of the simulated trajectories (Fig. 5F), representative participants were selected based on the relative stability of target items in the retained MCMC trajectories. For each selected participant, the final 300 retained MCMC sweeps were displayed as a binary item-by-iteration matrix, with rows corresponding to PHQ-9 items and columns corresponding to retained MCMC sweeps. A horizontal separator was placed between items 1–6 and items 7–9 to distinguish background and target items.

Flip rate was defined as

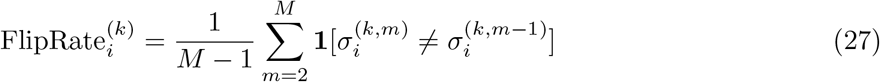

where **1**[·] is an indicator function that takes the value 1 when the condition is true and 0 when it is false. The flip-rate comparison was shown in Fig. 5G.

## Acknowledgements

We thank Yusuke Takahashi M.D. for helpful comments on the manuscript. This study was supported in part by the Moonshot R&D Program (JPMJMS2024-9 to H.N.) and CREST (JPMJCR25Q2 to H.N. and JPMJCR22N5 to T.A.K.) from the Japan Science and Technology Agency (JST), Japan Agency for Medical Research and Development (AMED) Multidisciplinary Frontier Brain and Neuroscience Discoveries (Brain/MINDS 2.0) (JP25wm0625322 to T.A.K. & H.N. and JP25wm0625210 to H.N.) and KAKENHI (21H03541 to H.N. and 22H00494 & 26H02458 to T.A.K.) from the JSPS, and Takeda Science Foundation (to M.T.).

## Author contributions

M.T.: Conceptualization, Methodology, Software, Formal analysis, Visualization, Writing – original draft, Writing – review & editing. H.N.: Conceptualization, Supervision, Writing – review & editing. T.K.: Data curation, Investigation, Resources, Writing – review & editing. T.A.K.: Data curation, Investigation, Resources, Writing – review & editing. All authors reviewed and approved the final manuscript.

## Data availability

The data that support the findings of this study are available from the corresponding author upon reasonable request.

## Code availability

The analysis code is available from the corresponding author upon reasonable request. This project is developed under Python 3.10.5 and is available on GitHub (https://github.com/masa10223/eladepression)

## Ethics Statement

This study was approved by the ethics committee of Kyushu University (25–84) and complied with all provisions of the Declaration of Helsinki.

