## appendix for "Mind the gap: quantifying individual–population gap in depressive symptom dynamics through energy landscapes"

### 1 Derivation of the VEM update equations

This appendix derives the variational expectation–maximization (VEM) updates from the evidence lower bound (ELBO). The starting point is

$$\mathcal{F} = \sum_{k=1}^{K} \mathbb{E}_{q^{(k)}}\left[\log p(\mathcal{D}^{(k)} \mid \boldsymbol{\theta}^{(k)})\right] - \sum_{k=1}^{K} \mathrm{KL}\Big(q^{(k)} \,\Big\|\, p(\boldsymbol{\theta}^{(k)} \mid \boldsymbol{\eta}, \boldsymbol{\alpha})\Big), \tag{17}$$

where

$$q^{(k)}(\boldsymbol{\theta}^{(k)}) = \mathcal{N}\Big(\boldsymbol{\theta}^{(k)} \mid \boldsymbol{\mu}^{(k)}, \mathrm{diag}(\boldsymbol{\beta}^{(k)})^{-1}\Big)$$

and

$$p(\boldsymbol{\theta}^{(k)} \mid \boldsymbol{\eta}, \boldsymbol{\alpha}) = \mathcal{N}\Big(\boldsymbol{\theta}^{(k)} \mid \boldsymbol{\eta}, \mathrm{diag}(\boldsymbol{\alpha})^{-1}\Big).$$

The log-likelihood for participant $k$ is

$$\log p(\mathcal{D}^{(k)} \mid \boldsymbol{\theta}^{(k)}) = T \left(\boldsymbol{\theta}^{(k)}\right)^{\top} \langle \tilde{\boldsymbol{\sigma}}^{(k)} \rangle_{\text{emp}} - T\psi(\boldsymbol{\theta}^{(k)}),$$

with

$$\langle \tilde{\boldsymbol{\sigma}}^{(k)} \rangle_{\text{emp}} = \frac{1}{T} \sum_{t=1}^{T} \tilde{\boldsymbol{\sigma}}^{(k)}(t).$$

The log-partition function is approximated by a second-order Taylor expansion around the current group mean $\boldsymbol{\eta}$:

$$\psi(\boldsymbol{\theta}^{(k)}) \approx \psi(\boldsymbol{\eta}) + \langle \tilde{\boldsymbol{\sigma}} \rangle_{\boldsymbol{\eta}}^{\top} \left(\boldsymbol{\theta}^{(k)} - \boldsymbol{\eta}\right) + \frac{1}{2} \left(\boldsymbol{\theta}^{(k)} - \boldsymbol{\eta}\right)^{\top} C_{\boldsymbol{\eta}} \left(\boldsymbol{\theta}^{(k)} - \boldsymbol{\eta}\right). \tag{18}$$

Here, $\langle \tilde{\boldsymbol{\sigma}} \rangle_{\boldsymbol{\eta}}$ is the model-expected sufficient-statistic vector under $\boldsymbol{\eta}$, and $C_{\boldsymbol{\eta}}$ is the covariance matrix of $\tilde{\boldsymbol{\sigma}}$ under $\boldsymbol{\eta}$. Let

$$\boldsymbol{c}_{\boldsymbol{\eta}} = (C_{\boldsymbol{\eta},11}, \ldots, C_{\boldsymbol{\eta},MM})^{\top}$$

denote the vector of diagonal elements of $C_{\boldsymbol{\eta}}$.

### 1.1 Expansion of the participant-specific ELBO

Using the definition of KL divergence,

$$-\text{KL}(q^{(k)} \| p) = \mathbb{E}_{q^{(k)}} \left[\log p(\boldsymbol{\theta}^{(k)} \mid \boldsymbol{\eta}, \boldsymbol{\alpha})\right] - \mathbb{E}_{q^{(k)}} \left[\log q^{(k)}(\boldsymbol{\theta}^{(k)})\right],$$

the contribution of participant $k$ to (17) can be written as

$$\mathcal{F}^{(k)} = \mathbb{E}_{q^{(k)}} \left[\log p(\mathcal{D}^{(k)} \mid \boldsymbol{\theta}^{(k)})\right] + \mathbb{E}_{q^{(k)}} \left[\log p(\boldsymbol{\theta}^{(k)} \mid \boldsymbol{\eta}, \boldsymbol{\alpha})\right] - \mathbb{E}_{q^{(k)}} \left[\log q^{(k)}(\boldsymbol{\theta}^{(k)})\right].$$

First, the likelihood term is

$$\mathbb{E}_{q^{(k)}} \left[\log p(\mathcal{D}^{(k)} \mid \boldsymbol{\theta}^{(k)})\right] = T \left(\boldsymbol{\mu}^{(k)}\right)^{\top} \langle \tilde{\boldsymbol{\sigma}}^{(k)} \rangle_{\text{emp}} - T\mathbb{E}_{q^{(k)}} \left[\psi(\boldsymbol{\theta}^{(k)})\right].$$

By (18),

$$\mathbb{E}_{q^{(k)}} \left[\psi(\boldsymbol{\theta}^{(k)})\right] \approx \psi(\boldsymbol{\eta}) + \langle \tilde{\boldsymbol{\sigma}} \rangle_{\boldsymbol{\eta}}^{\top} \left(\boldsymbol{\mu}^{(k)} - \boldsymbol{\eta}\right) + \frac{1}{2} \mathbb{E}_{q^{(k)}} \left[\left(\boldsymbol{\theta}^{(k)} - \boldsymbol{\eta}\right)^{\top} C_{\boldsymbol{\eta}} \left(\boldsymbol{\theta}^{(k)} - \boldsymbol{\eta}\right)\right].$$

For any random vector $\boldsymbol{x}$ and any symmetric matrix $A$,

$$\mathbb{E}[\boldsymbol{x}^{\top} A \boldsymbol{x}] = \mathbb{E}[\boldsymbol{x}]^{\top} A \mathbb{E}[\boldsymbol{x}] + \text{tr}\{A \, \text{Cov}(\boldsymbol{x})\}.$$

With $\boldsymbol{x} = \boldsymbol{\theta}^{(k)} - \boldsymbol{\eta}$, we have

$$\mathbb{E}_{q^{(k)}}[\boldsymbol{x}] = \boldsymbol{\mu}^{(k)} - \boldsymbol{\eta}, \qquad \text{Cov}_{q^{(k)}}(\boldsymbol{x}) = \text{diag}(\boldsymbol{\beta}^{(k)})^{-1}.$$

Therefore,

$$\begin{aligned} &\mathbb{E}_{q^{(k)}} \left[\left(\boldsymbol{\theta}^{(k)} - \boldsymbol{\eta}\right)^{\top} C_{\boldsymbol{\eta}} \left(\boldsymbol{\theta}^{(k)} - \boldsymbol{\eta}\right)\right] \\ &= \left(\boldsymbol{\mu}^{(k)} - \boldsymbol{\eta}\right)^{\top} C_{\boldsymbol{\eta}} \left(\boldsymbol{\mu}^{(k)} - \boldsymbol{\eta}\right) + \text{tr}\left[\text{diag}(\boldsymbol{\beta}^{(k)})^{-1} C_{\boldsymbol{\eta}}\right]. \end{aligned}$$

Thus, after substituting the Taylor approximation, the likelihood contribution is

$$\mathbb{E}_{q^{(k)}}\left[\log p(\mathcal{D}^{(k)} \mid \boldsymbol{\theta}^{(k)})\right] \approx T\left(\boldsymbol{\mu}^{(k)}\right)^{\top} \langle \tilde{\boldsymbol{\sigma}}^{(k)} \rangle_{\mathrm{emp}} - T\psi(\boldsymbol{\eta}) - T\langle \tilde{\boldsymbol{\sigma}} \rangle_{\boldsymbol{\eta}}^{\top}\left(\boldsymbol{\mu}^{(k)} - \boldsymbol{\eta}\right) - \frac{T}{2}\left(\boldsymbol{\mu}^{(k)} - \boldsymbol{\eta}\right)^{\top} C_{\boldsymbol{\eta}}\left(\boldsymbol{\mu}^{(k)} - \boldsymbol{\eta}\right) - \frac{T}{2}\operatorname{tr}\left[\operatorname{diag}(\boldsymbol{\beta}^{(k)})^{-1} C_{\boldsymbol{\eta}}\right].$$

Next, the Gaussian prior gives

$$\mathbb{E}_{q^{(k)}}\left[\log p(\boldsymbol{\theta}^{(k)} \mid \boldsymbol{\eta}, \boldsymbol{\alpha})\right] = \frac{1}{2}\sum_{j=1}^{M}\log\alpha_j - \frac{1}{2}\sum_{j=1}^{M}\alpha_j\left\{\left(\mu_j^{(k)} - \eta_j\right)^2 + \left(\beta_j^{(k)}\right)^{-1}\right\} + \text{const.}$$

Finally, the entropy term of the diagonal Gaussian variational distribution is

$$-\mathbb{E}_{q^{(k)}}\left[\log q^{(k)}(\boldsymbol{\theta}^{(k)})\right] = -\frac{1}{2}\sum_{j=1}^{M}\log\beta_j^{(k)} + \text{const.}$$

### 1.2 Derivation of the E-step update for the posterior mean

Collect only the terms depending on $\boldsymbol{\mu}^{(k)}$:

$$\mathcal{F}_{\mu}^{(k)} = T\left(\boldsymbol{\mu}^{(k)}\right)^{\top} \langle \tilde{\boldsymbol{\sigma}}^{(k)} \rangle_{\mathrm{emp}} - T\langle \tilde{\boldsymbol{\sigma}} \rangle_{\boldsymbol{\eta}}^{\top}\left(\boldsymbol{\mu}^{(k)} - \boldsymbol{\eta}\right) - \frac{T}{2}\left(\boldsymbol{\mu}^{(k)} - \boldsymbol{\eta}\right)^{\top} C_{\boldsymbol{\eta}}\left(\boldsymbol{\mu}^{(k)} - \boldsymbol{\eta}\right) - \frac{1}{2}\left(\boldsymbol{\mu}^{(k)} - \boldsymbol{\eta}\right)^{\top} \operatorname{diag}(\boldsymbol{\alpha})\left(\boldsymbol{\mu}^{(k)} - \boldsymbol{\eta}\right) + \text{const.}$$

Differentiating with respect to $\boldsymbol{\mu}^{(k)}$ yields

$$\frac{\partial \mathcal{F}_{\mu}^{(k)}}{\partial \boldsymbol{\mu}^{(k)}} = T\left(\langle \tilde{\boldsymbol{\sigma}}^{(k)} \rangle_{\mathrm{emp}} - \langle \tilde{\boldsymbol{\sigma}} \rangle_{\boldsymbol{\eta}}\right) - \left(TC_{\boldsymbol{\eta}} + \operatorname{diag}(\boldsymbol{\alpha})\right)\left(\boldsymbol{\mu}^{(k)} - \boldsymbol{\eta}\right).$$

Setting this derivative to zero gives

$$\left(TC_{\boldsymbol{\eta}} + \operatorname{diag}(\boldsymbol{\alpha})\right)\left(\boldsymbol{\mu}^{(k)} - \boldsymbol{\eta}\right) = T\left(\langle \tilde{\boldsymbol{\sigma}}^{(k)} \rangle_{\mathrm{emp}} - \langle \tilde{\boldsymbol{\sigma}} \rangle_{\boldsymbol{\eta}}\right).$$

Solving for $\boldsymbol{\mu}^{(k)}$ gives

$$\boldsymbol{\mu}^{(k)} = \boldsymbol{\eta} + \left\{TC_{\boldsymbol{\eta}} + \operatorname{diag}(\boldsymbol{\alpha})\right\}^{-1} T\left(\langle \tilde{\boldsymbol{\sigma}}^{(k)} \rangle_{\mathrm{emp}} - \langle \tilde{\boldsymbol{\sigma}} \rangle_{\boldsymbol{\eta}}\right). \tag{19}$$

### 1.3 Derivation of the E-step update for the posterior precision

Collect only the terms depending on $\boldsymbol{\beta}^{(k)}$:

$$\mathcal{F}_{\beta}^{(k)} = -\frac{T}{2}\operatorname{tr}\left[\operatorname{diag}(\boldsymbol{\beta}^{(k)})^{-1} C_{\boldsymbol{\eta}}\right] - \frac{1}{2}\operatorname{tr}\left[\operatorname{diag}(\boldsymbol{\beta}^{(k)})^{-1}\operatorname{diag}(\boldsymbol{\alpha})\right] - \frac{1}{2}\sum_{j=1}^{M}\log\beta_j^{(k)} + \text{const.}$$

Since $c_{\eta,j} = C_{\eta,jj}$,

$$\mathcal{F}_{\beta}^{(k)} = -\frac{T}{2}\sum_{j=1}^{M}\frac{c_{\eta,j}}{\beta_j^{(k)}} - \frac{1}{2}\sum_{j=1}^{M}\frac{\alpha_j}{\beta_j^{(k)}} - \frac{1}{2}\sum_{j=1}^{M}\log\beta_j^{(k)} + \text{const.}$$

Differentiating with respect to $\beta_j^{(k)}$ gives

$$\frac{\partial \mathcal{F}_\beta^{(k)}}{\partial \beta_j^{(k)}} = \frac{Tc_{\eta,j}}{2(\beta_j^{(k)})^2} + \frac{\alpha_j}{2(\beta_j^{(k)})^2} - \frac{1}{2\beta_j^{(k)}}.$$

Setting this derivative to zero and multiplying both sides by $2(\beta_j^{(k)})^2$ gives

$$Tc_{\eta,j} + \alpha_j - \beta_j^{(k)} = 0.$$

Therefore,

$$\boldsymbol{\beta}^{(k)} = T\boldsymbol{c}_{\boldsymbol{\eta}} + \boldsymbol{\alpha}. \tag{20}$$

### 1.4 Derivation of the M-step update for the group mean

In the M-step, the variational distributions $q^{(k)}$ are held fixed, and $\boldsymbol{\eta}$ and $\boldsymbol{\alpha}$ are updated. The terms depending on $\boldsymbol{\eta}$ and $\boldsymbol{\alpha}$ are

$$\mathcal{F}_{\eta,\alpha} = \frac{K}{2}\sum_{j=1}^{M} \log \alpha_j - \frac{1}{2}\sum_{k=1}^{K}\sum_{j=1}^{M} \alpha_j \left\{ \left(\mu_j^{(k)} - \eta_j\right)^2 + \left(\beta_j^{(k)}\right)^{-1} \right\} + \text{const.}$$

For a fixed component $j$, the $\eta_j$-dependent part is

$$\mathcal{F}_{\eta_j} = -\frac{1}{2}\alpha_j \sum_{k=1}^{K} \left(\mu_j^{(k)} - \eta_j\right)^2 + \text{const.}$$

Thus,

$$\frac{\partial \mathcal{F}_{\eta_j}}{\partial \eta_j} = \alpha_j \sum_{k=1}^{K} \left(\mu_j^{(k)} - \eta_j\right).$$

Setting this derivative to zero gives

$$\sum_{k=1}^{K} \left(\mu_j^{(k)} - \eta_j\right) = 0, \qquad \eta_j = \frac{1}{K}\sum_{k=1}^{K} \mu_j^{(k)}.$$

Therefore,

$$\boldsymbol{\eta} = \frac{1}{K}\sum_{k=1}^{K} \boldsymbol{\mu}^{(k)}. \tag{21}$$

### 1.5 Derivation of the M-step update for the group precision

For a fixed component $j$, collect the terms depending on $\alpha_j$:

$$\mathcal{F}_{\alpha_j} = \frac{K}{2}\log \alpha_j - \frac{1}{2}\alpha_j \sum_{k=1}^{K} \left\{ \left(\mu_j^{(k)} - \eta_j\right)^2 + \left(\beta_j^{(k)}\right)^{-1} \right\} + \text{const.}$$

Differentiating with respect to $\alpha_j$ gives

$$\frac{\partial \mathcal{F}_{\alpha_j}}{\partial \alpha_j} = \frac{K}{2\alpha_j} - \frac{1}{2}\sum_{k=1}^{K} \left\{ \left(\mu_j^{(k)} - \eta_j\right)^2 + \left(\beta_j^{(k)}\right)^{-1} \right\}. \tag{22}$$

Setting this derivative to zero yields

$$\alpha_j = \frac{K}{\sum_{k=1}^{K} \left\{ \left( \mu_j^{(k)} - \eta_j \right)^2 + \left( \beta_j^{(k)} \right)^{-1} \right\}}. \tag{23}$$

Equivalently, in componentwise vector notation,

$$\boldsymbol{\alpha} = K \oslash \sum_{k=1}^{K} \left[ \left( \boldsymbol{\mu}^{(k)} - \boldsymbol{\eta} \right) \odot \left( \boldsymbol{\mu}^{(k)} - \boldsymbol{\eta} \right) + \left( \boldsymbol{\beta}^{(k)} \right)^{-1} \right],$$

where $\oslash$ and $\odot$ denote componentwise division and multiplication, respectively.
